# Rac guanine nucleotide exchange factors promoting Lgl1 phosphorylation in glioblastoma

**DOI:** 10.1101/2020.12.01.406538

**Authors:** Sylvie J Lavictoire, Danny Jomaa, Alexander Gont, Karen Jardine, David P Cook, Ian AJ Lorimer

## Abstract

The protein Lgl1 has key roles in the regulation of cell polarity. We have shown that Lgl1 is inactivated by hyperphosphorylation in glioblastoma as a consequence of *PTEN* loss and aberrant activation of the PI 3-kinase pathway; this contributes to glioblastoma pathogenesis both by promoting invasion and repressing glioblastoma cell differentiation. Lgl1 is phosphorylated by atypical protein kinase C in a complex with Par6 and activated Rac. Here we have investigated the role of specific Rac guanine nucleotide exchange factors in Lgl1 hyperphosphorylation in glioblastoma. We used CRISPR/Cas9 to knockout PREX1, a PI 3-kinase pathway-responsive Rac guanine nucleotide exchange factor, in patient-derived glioblastoma cells. Knockout cells had reduced Lgl1 phosphorylation which could be reversed by re-expressing PREX1. PREX1 knockout cells showed reduced motility and an altered phenotype suggestive of partial neuronal differentiation; consistent with this, RNA-seq analyses identified sets of PREX1-regulated genes associated with changes in cell motility and neuronal differentiation. PREX1 knockout in glioblastoma cells from a second patient did not affect Lgl1 phosphorylation. These cells overexpressed a short isoform of the Rac guanine nucleotide exchange factor TIAM1; knockdown of TIAM1 in PREX1-knockout cells from this patient reduced Lgl1 phosphorylation. These data show that PREX1 links aberrant PI 3-kinase signaling to Lgl1 phosphorylation in glioblastoma, but that TIAM1 can also promote Lgl phosphorylation in a subset of patients. While this shows redundant mechanisms for Lgl1 phosphorylation, PREX1 appears to have a non-redundant role in glioblastoma cell motility, as this was impaired in PREX1 knockout cells from both patients.

## INTRODUCTION

Glioblastoma is an incurable form of brain cancer. *De novo* glioblastoma is the most common type of glioblastoma, accounting for approximately 90% of cases. While *de novo* glioblastoma exhibits extensive heterogeneity both at the histological and molecular levels, comprehensive genetic analyses show that aberrant activation of the PI 3-kinase pathway occurs in almost all patients (1,2). This occurs through partial or complete mutational inactivation of *PTEN*, amplification of the tyrosine kinase receptors *EGFR* or *PDGFRA* and/or activating mutations in PI 3-kinase. Mouse models show that central nervous system-specific *PTEN* haploinsufficiency, in a TP53-null background, generates a brain cancer that closely resembles human glioblastoma (3). Two key features that are present in the mouse and human disease are the highly invasive behaviour of the cancer and the presence of a population of cancer cells with a neural stem cell-like phenotype. With respect to the latter, *PTEN* inactivation, together with *TP53* inactivation, represses differentiation of cultured neural stem cells (4), suggesting a direct role for PTEN in maintaining and/or expanding a population of neural stem cell-like cells in glioblastoma.

The *Drosophila* mutant lethal 2 giant larvae was identified almost fifty years ago (5). This mutant shows overgrowth of brain tissue leading to death at the larval stage. Detailed studies have shown that the brain phenotype is due to a failure of neuroblasts to differentiate: rather than undergoing asymmetric cell divisions to produce a neuroblast and a committed neural progenitor, neuroblasts undergo symmetric divisions to produce two neuroblasts (6). Transplantation studies showed that these neuroblasts were also invasive within the *Drosophila* central nervous system (5). The lethal giant larvae phenotype is caused by deletion of the gene encoding the protein Lgl, a double beta-propeller protein with cytoskeletal protein-like functions that include binding to membranes and to non-muscle myosin II (7–10). Its activity is negatively regulated by phosphorylation, predominantly mediated by atypical protein kinase C (11). While initial studies suggested that phosphorylation caused a large conformational change in Lgl (12), recent crystallography studies have not supported this (7). Rather, these studies show that phosphorylation occurs on a surface loop that is rich in positively charged residues that mediate membrane association of Lgl. Phosphorylation directly counters membrane association by neutralizing this negatively charged region and also indirectly by preventing the lipid binding-induced formation of an alpha-helical segment within this loop that arranges positively-charged residues in a conformation that enhances their membrane interaction.

As mutational inactivation of Lgl in *Drosophila* causes a glioblastoma-like phenotype, we asked whether Lgl inactivation might also have a role in human glioblastoma. In humans there are two genes encoding homologs of *Drosophila* Lgl, *LLGL1* and *LLGL2*. Neither of these is mutated in glioblastoma and *LLGL1*, encoding the protein Lgl1, is expressed relatively abundantly. We explored the possibility that Lgl1 was instead inactivated by hyperphosphorylation. Consistent with this, we showed that *PTEN*-null glioblastoma cells had a high level of Lgl1 phosphorylating activity and that this was reduced upon restoration of PTEN expression (13). Introduction of a non-phosphorylatable, constitutively-active version of Lgl1 repressed glioblastoma cell invasion and promoted its differentiation, both in cell culture and *in vivo* (14).

Lgl binds the scaffolding protein Par6; Par6 also binds aPKC and it is this complex that mediates Lgl phosphorylation (15). Activation of aPKC is controlled by binding of a third protein to Par6, either activated (*i.e.* GTP-bound) Cdc42 or Rac GTPases (16). Of the three Par6 protein family members (*PARD6A*, *PARD6B* and *PARD6G*), TCGA RNA-Seq data suggests that *PARD6A* is the most highly expressed (1). Par6A, encoded by *PARD6A*, is able to bind both Cdc42 and Rac (16). Pulldowns of flag-tagged Par6A in glioblastoma cells showed that it predominantly associated with Rac1 (17). This association requires Rac1 activation by a Rac guanine nucleotide exchange factor (GEF). Here we have assessed the role of specific Rac GEFs in promoting the phosphorylation of Lgl1 in glioblastoma cells isolated from patients. We initially focussed on the Rac GEF PREX1, which we previously showed was overexpressed in glioblastoma relative to normal brain (17). CRISPR/Cas9 knockout of PREX1 in glioblastoma cells from one patient showed markedly reduced Lgl1 phosphorylation, along with reduced motility and an apparent partial differentiation along the neuronal lineage. Knockout of PREX1 in cells from a second patient did not affect Lgl1 phosphorylation; these cells overexpressed a second Rac GEF, TIAM1, which was able to promote Lgl1 phosphorylation in the absence of PREX1, showing that there are redundant mechanisms for Lgl1 phosphorylation in a subset of glioblastoma patients. However, knockout of PREX1 in cells from both patients reduced motility, suggesting a non-redundant role for PREX1 in driving glioblastoma invasion.

## RESULTS

### Generation of PriGO8A PREX1 knockout cells

PriGO8A cells are glioblastoma cells that were isolated from a patient undergoing surgery for glioblastoma. They were isolated and cultured in neural stem cell media using monolayer growth on laminin substrate and a 5 % oxygen environment (13). Details of their neural stem cell-like properties and *in vivo* growth after intracerebral injection into immunocompromised mice have been described previously (13,14). To knock out PREX1 in PriGO8A cells, complexes of Cas9, tracrRNA and a crRNA targeting exon 2 of PREX1 were electroporated into cells as described in Materials and Methods. To produce a high frequency of mutated alleles, the electroporation was repeated two weeks later. Clonal populations were isolated using limiting dilution, with 29 wells in a 96 well plate giving cell populations. PriGO8A cells grow poorly at low densities: to compensate for this, clonal populations were isolated using conditioned media prepared as described in Materials and Methods. The mutation status of clones was assessed by TIDE analysis (18). The majority of isolated clones had a mixture of −5 and −2, consistent with clonal populations with biallelic PREX1 mutations. 8A/clone 4, along with a second clone with the same pattern of *PREX1* mutations (8A/clone 6), were chosen for further study (Figure 1A). An additional population (8A/clone 18) had −5 and −2 deletions and +1 insertions (Figure 1A). Although this is referred to as clone 18 here, this may be a mixed population of cells with different *PREX1* mutations. This was also chosen for initial further study as it lacked detectable wild type *PREX1* alleles. Western blot analysis showed that the three clones did not express detectable PREX1 protein (Figure 1B) and immunofluorescence analysis of 8A/clone 4 showed that PREX1 protein expression was uniformly absent from individual cells (Figure 1C).

**Figure 1.**
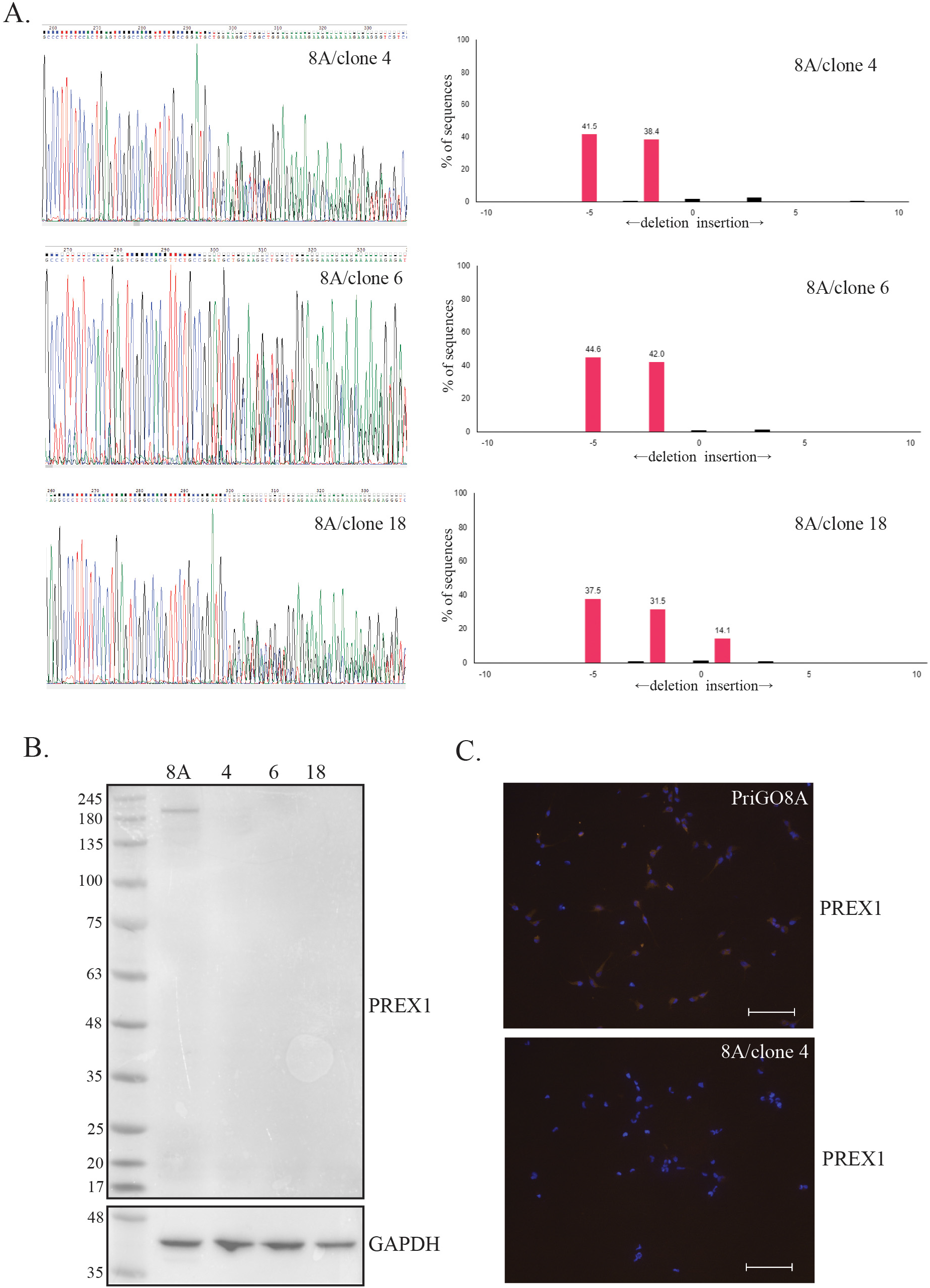
Generation of PREX1 knockout PriGO8A patient-derived glioblastoma cells. **A**. Genomic analysis of PREX1 knockout cells. To determine deletions/insertions in clones isolated by limiting dilution, the targeted region of the *PREX1* gene was PCR amplified and sequenced (left panels). Sequence trace decomposition (right panels) was done using software described in Brinkman *et al*.(29). **B**. Western blot showing PREX1 expression in parental cells (8A) and three knockout clones. **C**. Immunofluorescence for PREX1 on PriGO8A cells and clone 4. Scale bar is 100 um.

### Lgl1phosphorylation in PREX1-null glioblastoma cells

The ability of PREX1-null clones to phosphorylate Lgl1 was compared to parental PriGO8A cells. As antibodies that detect endogenous Lgl1 phosphorylation are not available, PriGO8A cells were first transduced with flag-tagged Lgl to increase levels. Total cell lysates were then probed by Western blotting with an antibody that detects phosphorylated PKC substrates. As described previously, this detects a band of the expected size for Lgl1 (kDa) that is absent in untransduced cells and cells transduced with a version of Lgl1 in which the three hinge region serine residues are mutated to alanine (13); the band is also reduced by siRNA knockdown of Lgl1 and atypical PKCɩ (13). In the two PREX1-null clones examined here, levels of phosphorylated Lgl1 were consistently lower compared to PriGO8A parental cells (Figure 2A). Levels of total transduced Lgl1 protein were also consistently lower, but quantitation showed that the overall extent of Lgl1 phosphorylation was still lower in the PREX1-null clones when this was corrected for (Figure 2A).

**Figure 2.**
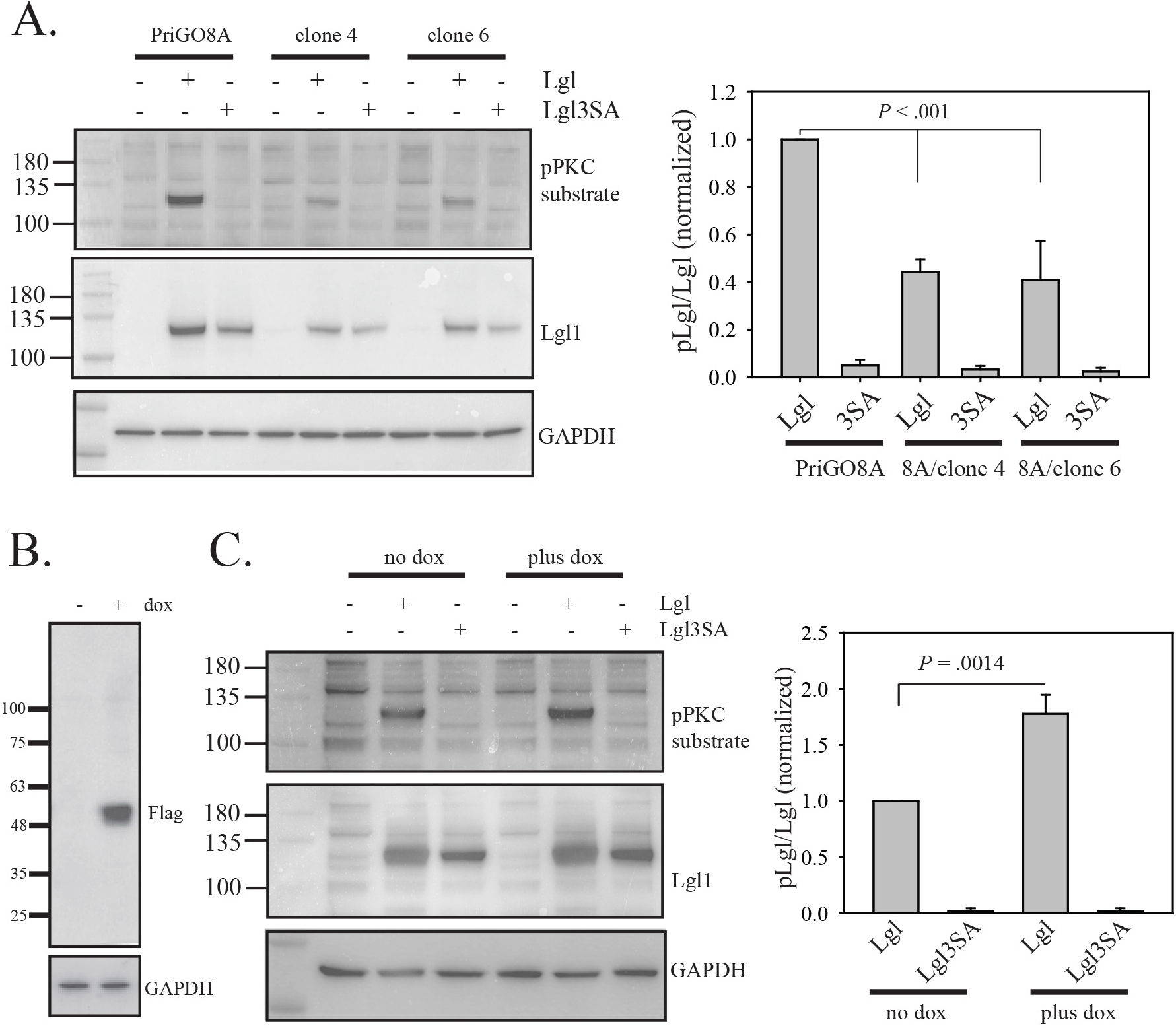
Lgl1 phosphorylation in PriGO8A and PREX1-knockout cells. **A**. PriGO8A cells and PREX1-null knockout clones 4 and 6 were either untransduced, transduced with lentivirus expressing Lgl, or transduced with lentivirus expressing Lgl3SA. Two days later, samples were collected for Western blotting with antibody to phosphoPKC substrate, Lgl and GAPDH. **B**. Induction of Flag-tagged PREX PHDH domain expression in PREX1-null cells. Transduced clone 4 cells were either untreated or treated for 48 with 1 μg/ml doxycycline. Cells were harvested and analyzed by Western blotting with anti-Flag antibody. **C.** PREX1-knockout cells transduced with lentiviral vectors for doxycycline-inducible expression of PREX1 were plated in the absence or presence of doxycycline. One day later, cells were either untransduced, transduced with lentivirus expressing Lgl, or transduced with lentivirus expressing Lgl3SA. Two days later, samples were collected for Western blotting with antibody to phosphoPKC substrate, Lgl, and GAPDH. For A and B, bar graphs show quantitative data from three different experiments. Data for each experiment were normalized to the pLgl/Lgl signal for cells transduced with Lgl (for B, in the absence of doxycycline). *P* values were determined by One Way Analysis of Variance/ All Pairwise Multiple Comparison Procedures (Holm-Sidak method) in A and with a two-tailed t test in B.

Differences in Lgl1 phosphorylation could be explained by loss of PREX1 but could alternatively be due off-target effects of the crRNA, clonal variation, or changes due to the long-term passage of these cells that was necessary to isolate clonal populations. To determine if the changes were due to loss of PREX1, PREX1-null glioblastoma cells were genetically modified for doxycycline-inducible expression of PREX1 (Figure 2B). To facilitate the production of high titre lentivirus, lentivirus expressing only the DHPH domain of PREX1 (with a carboxy-terminal Flag tag) was made. The DHPH domain of PREX1 contains the Rac1 guanine nucleotide exchange factor activity and also retains its regulation by PIP3 and G protein βγsubunit binding(19). The reduced ability of PREX1-null clone 4 cells to phosphorylate Lgl1 was reversed by induction of PREX1 expression with doxycycline, confirming that this effect was due to the knockout (Figure 2C).

### Morphology motility and differentiation state of PREX1null cells

Phase contrast video-microscopy of multiple PREX1-null clones showed that they had an altered morphology with a smaller cell body and numerous thin branched extensions that extended and retracted dynamically (Figure 3A and Supporting information videos S1-4). This morphology became apparent approximately two days after plating cells. PREX1-null cells also did not form lamellipodia, which are readily detected in parental cells. Induction of PREX1 reversed the morphological changes seen in PREX1-null cells, with cells showing fewer neurite-like extensions, a larger cell body, and abundant lamellipodia (Figure 3B and Supporting information videos S5 and S6). PREX1-null glioblastoma cells have reduced motility, as expected based on our earlier study (17) (Figure 4A and Supporting information videos S1-4). Induction of PREX1 reversed this loss of motility, consistent with the appearance of lamellipodia ((Figure 4B and Supporting information videos S5 and S6)).

**Figure 3.**
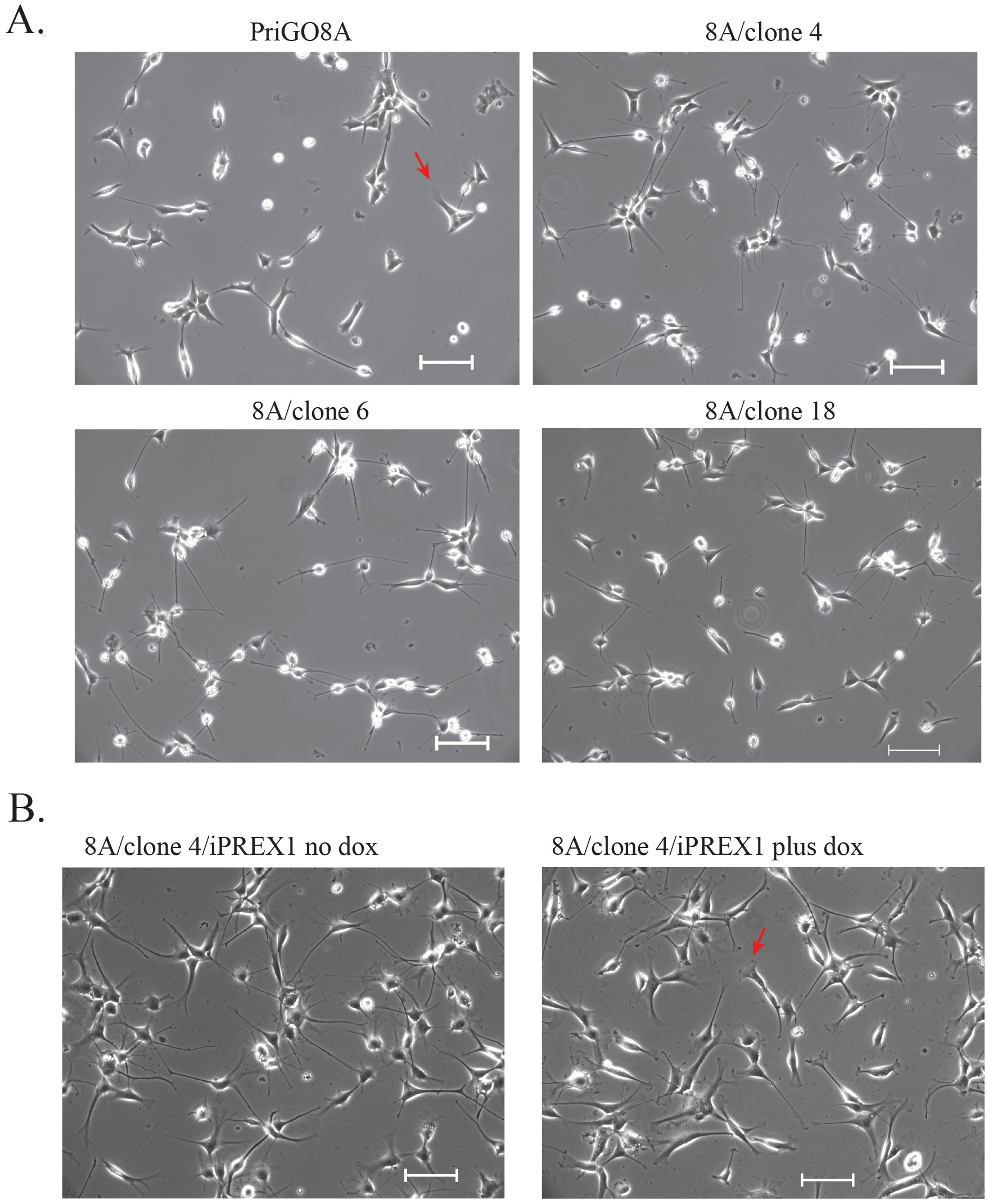
Morphology of PREX1-knockout PriGO8A cells. **A.** Live cell phase contrast images of PriGO8A and clones 4, 6 and 18. **B.** Live cell phase contrast images of PriGO8A PREX1-knockout cells transduced with lentiviral vectors for doxycycline-inducible PREX1 and grown in the absence of presence of doxycycline. Scale bars are 200 μm. Red arrows indicate examples of lamellipodia. Full microscopy videos are shown in the supplementary data videos S1-S6.

**Figure 4.**
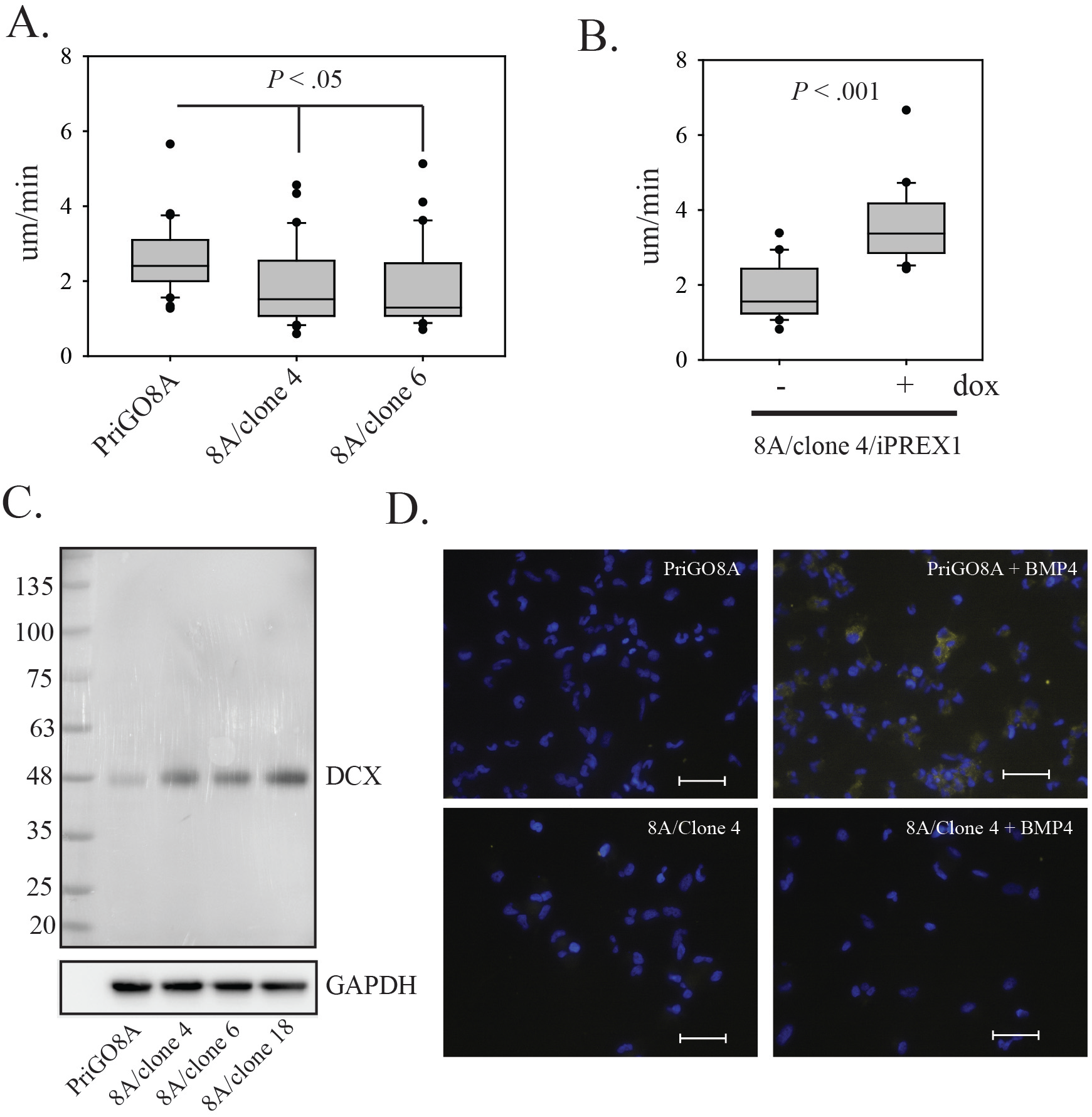
Changes in motility and differentiation state in PriGO8A PREX1-knockout cells. **A** and **B.** Motility of PREX1-null clones 4 and 6 (A) and PREX1-null clone 4 cells with re-expression of PREX1. The Y axis shows rate of movement in um/min. Box plots show the mean and 25^th^ and 75^th^ percentiles, with whiskers showing the 10^th^ and 90^th^ percentiles. The *P* value for A was determined using the Kruskal Wallis One Way Analysis on Ranks/All Pairwise Multiple Comparison Procedures (Tukey Test). The *P* value for B was determined using the Mann-Whitney Rank Sum test. See also supplementary videos. **C.** Total cell lysates from PriGO8A cells and three knockout clones were analyzed for expression of doublecortin by Western blot analysis. **D.** PriGO8A and 8A/clone 4 cells were treated with 100 ng/ml BMP4 for 6-7 days. Cells were then fixed and immunofluorescence for GFAP was performed. Scale bar is 100 μm.

As the morphology of PREX1-null clones was suggestive of partial differentiation along the neuronal lineage, the expression of doublecortin, a marker of committed neural progenitors/immature neurons (20), was assessed. Western blot analysis showed that PREX1 knockout cells expressed increased levels of doublecortin (Figure 4C). Consistent with data from other glioblastoma cells isolated under serum-free conditions (21), PriGO8A cells undergo differentiation along the astrocytic, but not the neuronal, lineage when treated with BMP4 (22). To determine if PREX1-null glioblastoma cells had lost their multilineage potential, we treated with them with BMP4. Although control PriGO8A cells became positive for GFAP expression as expected, PREX1 null cells did not. (Figure 4D). The morphology changes, doublecortin expression and loss of ability to differentiate along the astrocytic lineage, are all consistent PREX1-null cells having undergone partial neuronal differentiation.

### RNA-seq analysis of PREX1 knockout cells

To further characterize PREX1-null glioblastoma cells, RNA-seq analyses were performed, comparing parental PriGO8A cells with 8A/clone 4 cells and also comparing 8A/clone 4 cells with doxycycline-inducible PREX1, without and with 48 h doxycycline treatment. Each analysis was performed on two biological replicates. Figure 5A shows the number of genes with significantly changed expression in the two analyses at different cutoffs. In the PriGO8A:8A/clone 4 comparison, there were a large number of gene changes, with more genes down-regulated than upregulated. In the comparison of 8A/clone 4/inducible PREX1 cells without and with doxycycline treatment, a much smaller number of genes showed changed expression, with upregulated genes predominating. These patterns are consistent with clone 4 cells having undergone partial differentiation (large number of expression changes, with down-regulation predominating due to cell specialization) and short-term activation of a specific signaling pathway in clone 4/DHPH treated with doxycycline. PREX1 mRNA levels were significantly lower in clone 4 (b = −0.55), potentially due to either changes in transcription or nonsense-mediated decay. Recent work has shown that gene knockouts can have compensatory increases in transcription of related genes by a mechanism that involves nonsense-mediated decay (23). However, changes in PREX2 mRNA, the gene most closely related to PREX1, were not observed, showing that this compensatory mechanism is not active in the PREX1 knockouts. With the inducible system, after 48 h treatment with doxycycline PREX1 mRNA expression was three-fold higher than in parental PriGO8A cells.

**Figure 5.**
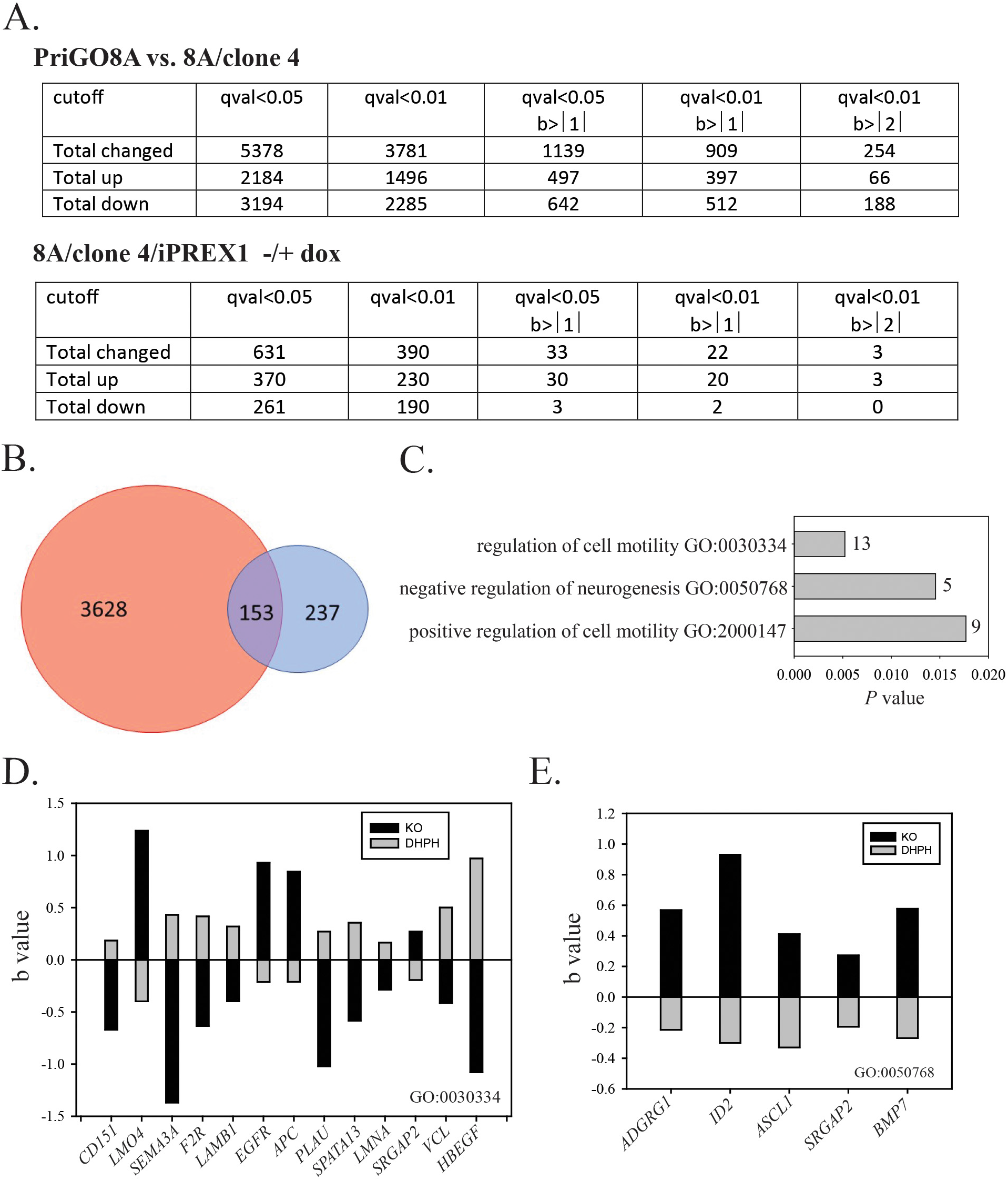
Changes in gene expression in PREX1 knockout cells. **A.** Number of changes in gene expression using various cut-offs. q values (qval) are p values adjusted for multiple comparisons using the Benjamini-Hochberg FDR method. b values are estimates of effect size based on beta coefficients of the linear model fit to each gene and are expressed in log2. **B**. Venn diagram using qval<0.01 data for both RNA-seq experiments. Red circle shows the 3781 genes with altered expression at this cutoff in the comparison of PriGO8A cells with clone 4 PREX1 knockout cells. Blue circle shows the 390 genes with altered expression in clone 4 cells engineered for doxycycline-inducible PREX1, with and without 48 h doxycycline treatment. 153 genes show altered expression in both data sets. **C.** Analysis of gene ontology biological processes for 153 gene overlap using Enrichr^7^. GO terms are shown on the y axis and *P* values on the x axis. Numbers of genes associated with the GO terms are shown beside the bars. **D** and **E.** Genes associated with GO terms for motility (D) and neurogenesis (E). Black bars show the change in expression between PriGO8A and 8A/clone 4 cells; gray bars show the change in expression in 8A/clone 4/iPREX1 cells without and with 48 induction of PREX1 expression with doxycycline.

Figure 5B shows the intersection of the data sets from the RNA-seq analyses for PREX1 and knockout and restoration, using the qval <0.01 cutoff for both. This identified a set of 153 genes that are significantly altered (*i.e.* both increased and decreased expression) in both analyses. This set should identify genes which are regulated relatively directly by PREX1, although it will exclude PREX1-regulated genes where expression is only transient due to feedback inhibition or other factors. The set will also exclude differences due to clonal selection of clone 4 from the parental PriGO8A population and possible non-specific effects of doxycycline on gene expression. Analysis of the 153 gene set for GO biological processes using Enrichr gave three matches that were significant by Enrichr-assigned adjusted *P* values (excluding matches with less than five genes). These were “regulation of cell migration”, “negative regulation of neurogenesis” and “positive regulation of cell motility” (Figure 5C - E). Although the number of genes associated with negative regulation of neurogenesis was small, it notably included *ASCL1,* which has a well-known role in promoting neurogenesis (24).

### PREX1 knockout in glioblastoma cells from a second patient (PriGO9A)

For comparison, PREX1-null cells were generated in glioblastoma cells from a second patient, PriGO9A cells. The basic properties of PriGO9A cells were described previously (13). Six clones generated by limiting dilution were screened. Five of the six showed weak expression of PREX1 by Western blot; TIDE analysis showed that the majority of these had biallelic mutations of PREX1, but also a significant signal for wild type PREX1. One clone was identified that had no signal by Western blot and had triallelic mutations for *PREX1* (1, 2 and 4 base pair deletions, Figure 6A). Based on this, PriGO9A cells likely have three copies of the *PREX1* gene. This occurs in 37% of glioblastomas as a result of gain of an entire copy of chromosome 20 (25). In contrast to the findings with PREX1 knockouts in PriGO8A cells, PREX1-null PriGO9A cells showed no change in their ability to phosphorylate Lgl1 (Figure 6B) and did not show an obvious change in morphology (Figure 7A and Supporting information videos S7-S10). They did however show reduced motility that was restored upon re-expression of PREX1 (Figure 7B); videomicroscopy showed that while they extend small lamellipodia, these retract without productive cell movement.

**Figure 6.**
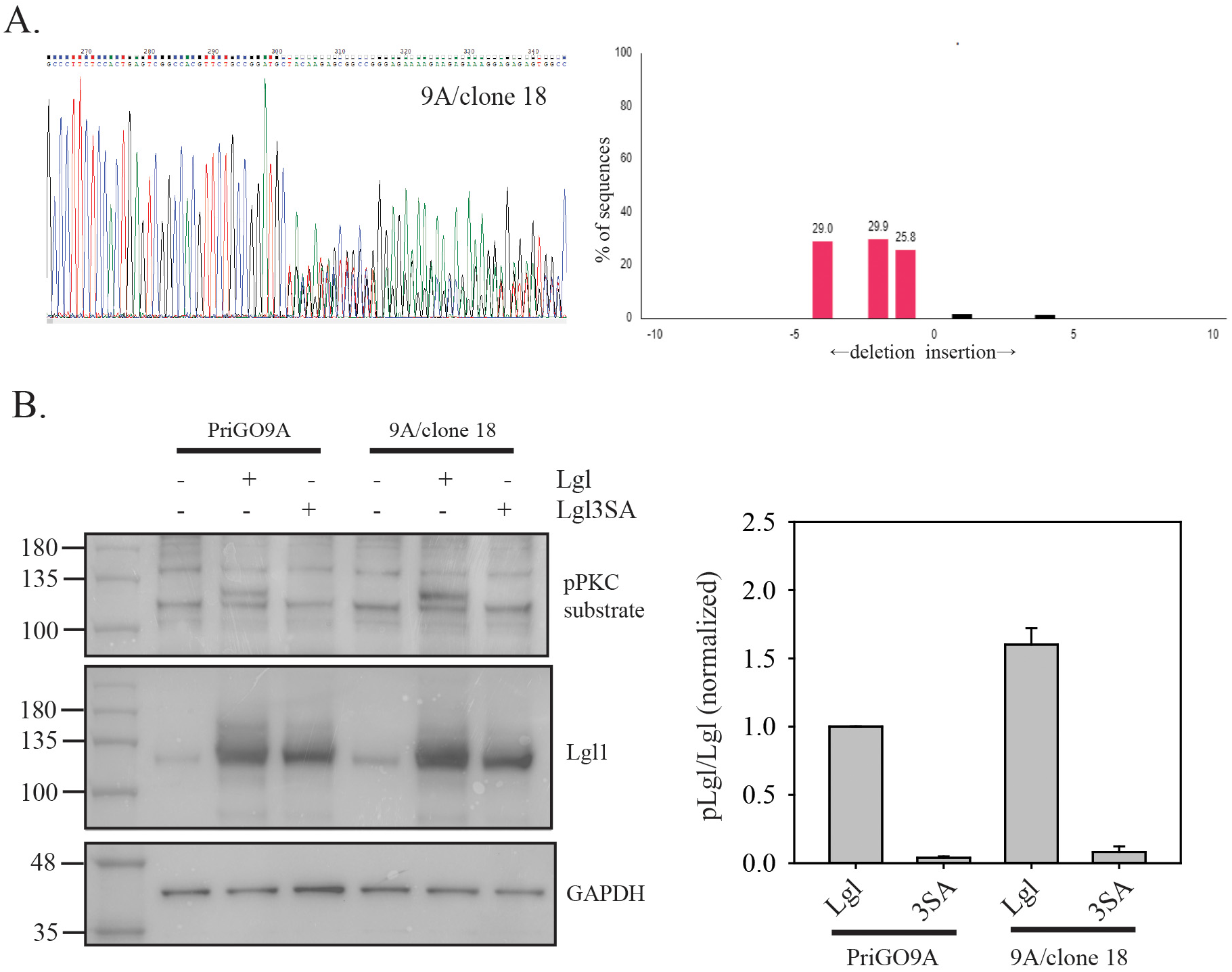
Generation of PREX1 knockout PriGO9A patient-derived glioblastoma cells. **A.** Genomic analysis of PriGO9A PREX1 knockout cells was performed as described in Figure 1A. **B.** Lgl phosphorylation in PriGO9A and PriGO9A PREX1 knockout cells was preformed as described in Figure 2A. The bar graph shows quantitative data from two separate experiments. Error bars shows the range. Data for each experiment were normalized to the pLgl/Lgl signal for PriGO9A cells transduced with Lgl.

**Figure 7.**
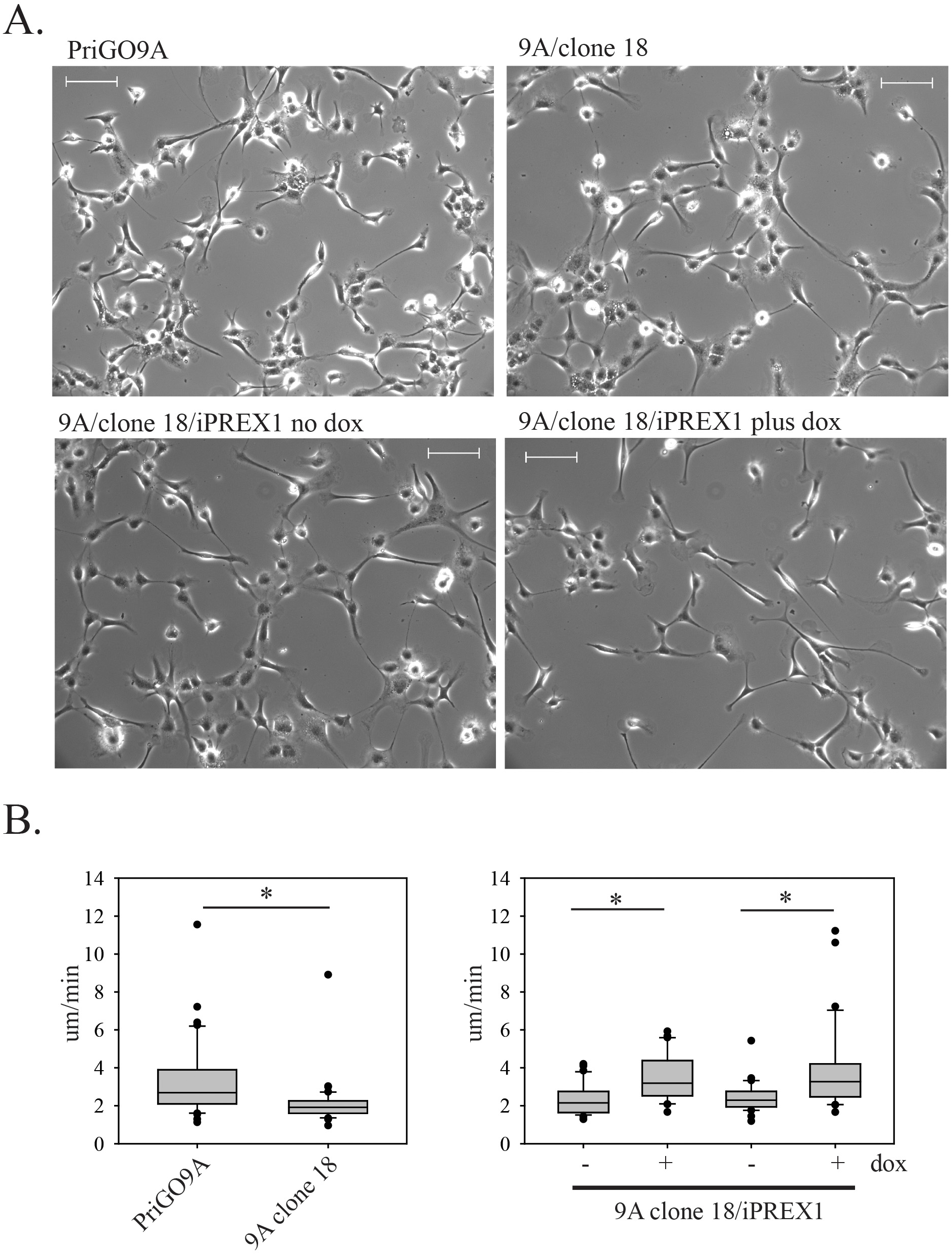
Morphology of PREX1-knockout PriGO9A cells. **A.** Top panels show live cell phase contrast images of PriGO9A (WT) and clone 18 (KO). Bottom panels show clone 18 cells transduced with lentiviral vectors for doxycycline-inducible expression of PREX1 and grown in the absence or presence of doxycycline. Scale bars are 200 μm. Full microscopy videos are shown in the supplementary data videos S7-S10 **B.** The left bar graph shows the motility of PriGO9A and 9A clone 18 cells, while the right panel show replicate experiments of 9A clone 18 cells with doxycycline-inducible PREX1. For both graphs, the Y axis shows rate of movement in um/min. Box plots show the mean and 25^th^ and 75^th^ percentiles, with whiskers showing the 10^th^ and 90^th^ percentiles. *P* values were determined using the Mann-Whitney Rank Sum test. See also supplementary videos.

One explanation for the differences between PriGO8A and PriGO9A PREX1 knockouts is that PriGO9A expressed high levels of a second Rac GEF that can also promote Lgl1 phosphorylation. Analysis of microarray expression data from PriGO8A, PriGO9A and two other patient-derived cultures(22) showed that PriGO9A cells expressed 5-12 fold higher levels of TIAM1 mRNA compared to cells from other patients (Figure 8A). TIAM1 protein was also expressed at higher levels (Figure 8B). Full-length TIAM1 is 1591 amino acids, with a predicted molecular weight of 178 kD. There is a small increase in a band of approximately this size, but most of the increase is in two smaller forms. As the antibody used recognizes a carboxy terminal region, these are likely amino terminal-truncated versions. Two of these, arising through alternate splicing, have been described (NP_001340613.1 and NP_001340614.1) with predicted molecular weights of 71 and 68 kD. Analysis of TCGA data using TCGASpliceSeq (26) indicates that glioblastomas more frequently express codons 18-29 of TIAM1, consistent with generation of these alternately-spliced versions (Figure 8C). TIAM1 has previously been shown to control activation of the Par polarity complex in other contexts (27). Analysis across 152 patients from the TCGA database suggests that PREX1 mRNA expression is much higher than TIAM1 in most patients (Figure 8D).

**Figure 8.**
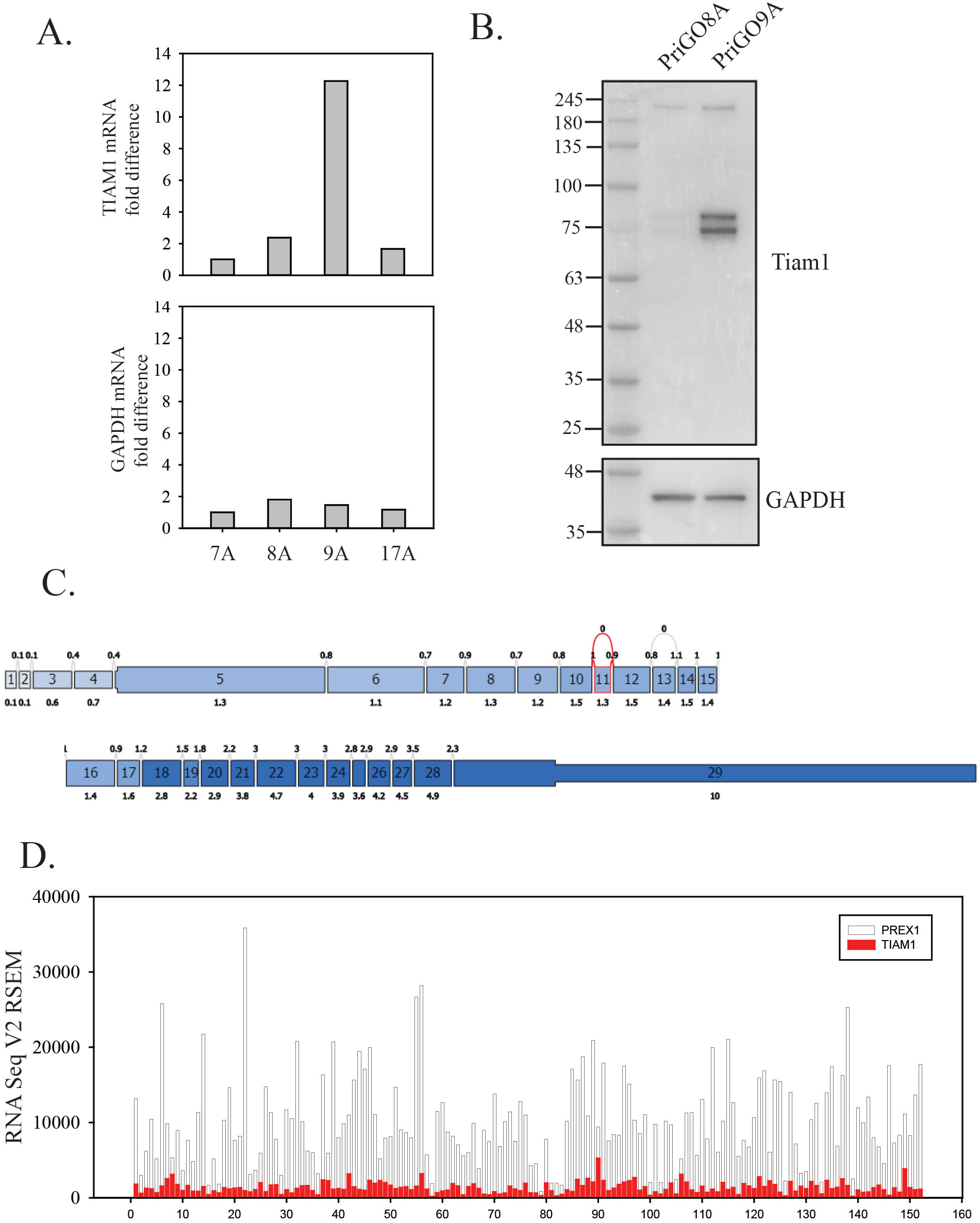
TIAM1 expression in glioblastoma. **A.** Relative mRNA expression in glioblastoma cells from four patients (PriGO7A, PriGO8A, PriGO9A and PriGO17A). Data are normalized to expression in PriGO7A. **B.** TIAM1 protein expression in PriGO8A and PriGO9A cells. Total cell lysates were analyzed by Western blotting. **C.** Analysis of TIAM1 slice variants in glioblastoma from TCGA RNA-seq data. Data analysis and figure generation were done with TCGA SpliceSeq (https://bioinformatics.mdanderson.org/public-software/tcgaspliceseq/). Numbered boxes show individual exons. Upper numbers show normalized read counts for splice events; lower numbers show normalized read counts for exon usage. The shading indicates expression level, with darker blue corresponding to higher expression. **D.** mRNA expression of PREX1 and TIAM1 from TCGA RNA-seq data. Y axis shows normalized read counts. X axis shows individual patients (152 in total). Clear bars and red bars show PREX1 and TIAM1 expression, respectively.

To determine if high TIAM1 expression was maintaining Lgl1 phosphorylation in the PriGO9A PREX1 knockout cells (9A/clone 18), RNA interference was used to deplete these cells of TIAM1. Initially a pool of siRNAs was used, which gave knockdown of the three major TIAM1 species. For further experiments, an siRNA from this pool that targets a 3’ region shared by common TIAM1 splice variants was used. This also gave knockdown of the three major TIAM1 species (Figure 9A). TIAM1 knockdown significantly reduced Lgl1 phosphorylation in these cells (Figure 9B and C). As a control, the effects of knockdown in PriGO8A cells was also assayed. TIAM1 knockdown in these cells had no effect on Lgl1 phosphorylation (Figure 9D).

**Figure 9.**
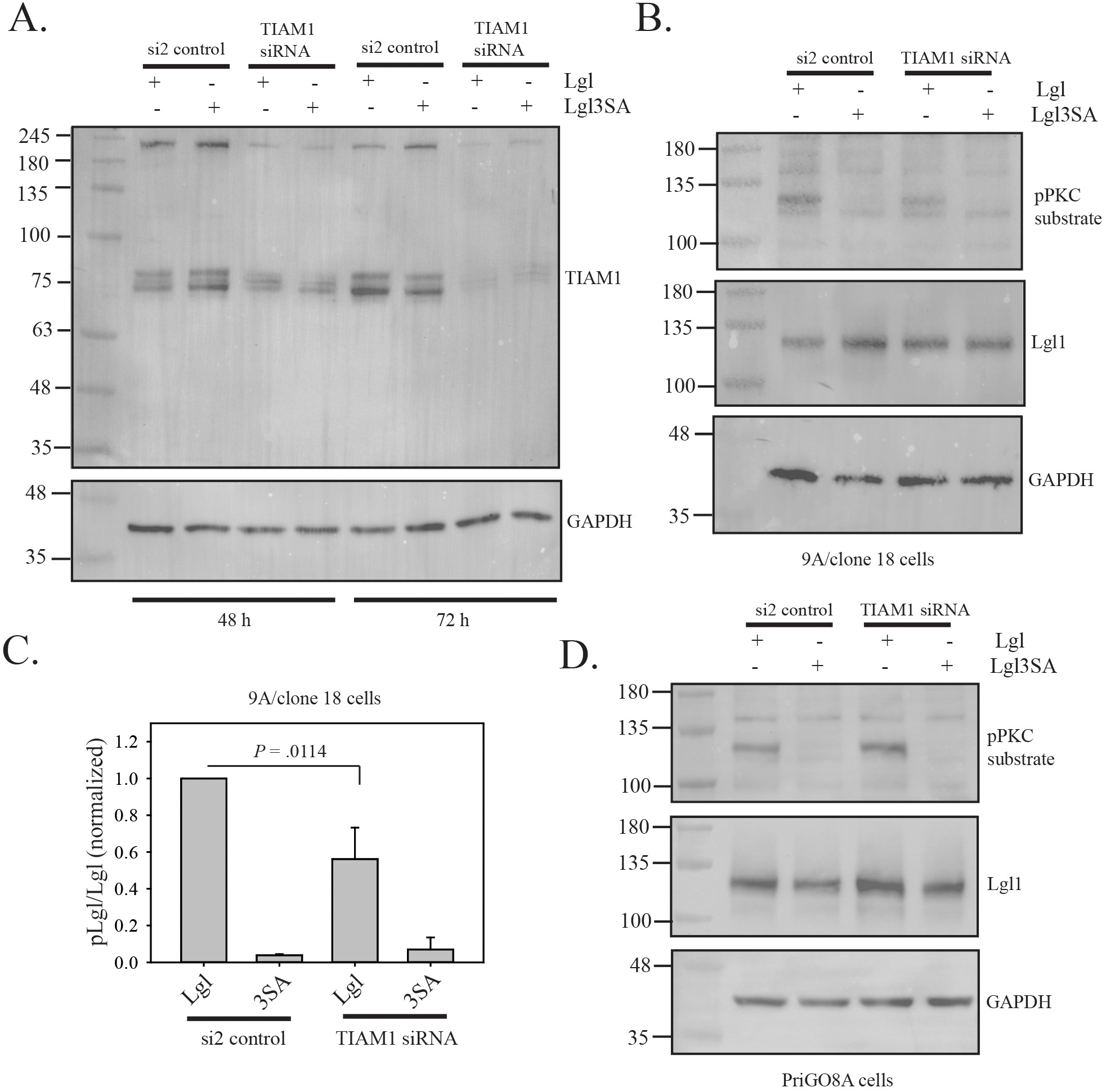
Effects of TIAM1 knockdown on Lgl phosphorylation. **A.** PriGO9A/clone 18 cells were transfected with a control siRNA (si2) or siRNA targeting TIAM1 one day after plating. One or two days after plating, cells were transduced with lentiviral vectors expressing either Lgl or Lgl3SA. Three and four days after plating (48 or 72 h post siRNA transfection), total cell lysates were collected and analyzed by Western blotting for TIAM1. **B.** PriGO9A/clone 18 cells were treated as in A. 72 h post siRNA transfection, total cell lysates were collected and analyzed by Western blotting for phosphoPKC substrate and Lgl. **C.** Bar graph bar showing quantitative data from three different experiments performed as in B. Data were normalized as describe in Figure 2. Error bars show standard deviation. The *P* value was determined using the Mann-Whitney Rank Sum Test. **D**. The same experiment as shown in B was performed, except that PriGO8A cells were used.

## DISCUSSION

Our previous work showed that inactivation of Lgl by phosphorylation has an important role in glioblastoma pathogenesis, where it is required both for the promotion of invasion and the repression of glioblastoma cell differentiation (14). Lgl phosphorylation was repressed when PTEN expression was restored in PTEN-null glioblastoma cells, showing a link between a common glioblastoma mutation and Lgl regulation (13). Here we have explored the role of Rac guanine nucleotide exchange factors in mediating signalling between the PI 3-kinase pathway and Lgl. We initially focussed on PREX1, as it is directly regulated by PIP3 binding(28) and is overexpressed in glioblastoma (17). We used CRISPR/Cas9 to knockout PREX1 in PriGO8A glioblastoma cells. PriGO8A cells were isolated from a glioblastoma patient under conditions that preserve neural stem cell-like characteristics and are *PTEN*-null, C250T *TERT* promoter mutant and without *EGFR* amplification (13). We were able to isolate clones that were confirmed to be PREX1-null by TIDE sequence analysis (18,29) and Western blotting, showing that PREX1 is not essential for glioblastoma cell growth in cell culture. To determine Lgl phosphorylating activity in these cells, we transduced them with Lgl and then detected phosphorylation using an antibody to phosphoPKC substrate as described previously (13). Transduction in parallel with a non-phosphorylatable version of Lgl serves as a control in these assays. Lgl1 phosphorylation was significantly reduced in PREX1 knockout cells and could be restored by re-expressing PREX1, confirming that this effect was directly due to the knockout. This identifies a novel role for PREX1 as a regulator of polarity.

Live cell imaging showed that PREX1 knockout cells no longer formed lamellipodia and had reduced motility, consistent with previous PREX1 siRNA knockdown experiments. Both lamellipodia formation and motility were restored with induction of PREX1 expression, confirming that the alterations seen in knockout cells were due to PREX1 loss. Live cell imaging also showed that multiple PREX1 clones had an altered morphology, with a smaller cell body and increased formation of thin, branched neurite-like extensions. This was suggestive of partial differentiation along the neuronal lineage. Consistent with this, knockout cells showed increased expression of the doublecortin, an early marker of neuronal differentiation; in addition, knockout cells no longer underwent astrocytic differentiation in response to BMP4 treatment, consistent with loss of multipotency. While some morphological changes are reversible, we have not fully evaluated the extent to which this partial differentiation is reversible, which will require a detailed epigenetic analysis. To further characterize PREX1 knockout cells, RNA-seq analysis of PriGO8A and 8a/clone 4 cells, as well as 8A/clone 4 cells with doxycycline-inducible PREX1, was performed. Analysis of the set of genes with changes in expression in both the PriGO8A: 8A/clone 4 comparison and the 8A/clone 4 without and with PREX1 induction gave gene ontology signatures corresponding to regulation of cell motility and neurogenesis, consistent with the conclusions from the microscopy and marker expression analyses. For the motility signature, *HBEGF* is notable as its positive regulation by PREX1 potentially creates a positive feed-back loop that could mediate a non-motile/motile switch. Notable in the negative regulation of neurogenesis signature is *ASCL1*, which is known to undergo a switch from an oscillating to sustained expression during neural stem cell differentiation into committed progenitors (30).

To determine how generalizable the above findings were, we also generated PREX1 knockouts in glioblastoma cells from a second patient (PriGO9A cells). While PriGO8A cells appears to have two copies of PREX1, based on TIDE analysis PriGO9A cells appear to have three copies, given that the one clone we isolated had three different deletions of roughly equal abundance. In contrast to PriGO8A PREX1 knockout cells, PriGO9A knockout cells did not show reduced Lgl phosphorylation, were still able to extend lamellipodia and did not show morphological features suggestive of differentiation along the neuronal lineage. They did, however, show reduced motility that could be restored by re-expressing PREX1. We reasoned that the observed differences might be due to expression of another Rac GEF in these cells with partially overlapping functions. Inspection of microarray data on these cells (22) showed that they expressed much higher mRNA levels of the Rac GEF TIAM1 and Western blotting confirmed that TIAM1 protein was also expressed at high levels in these cells. TIAM1 has previously been linked to polarity regulation in keratinocyte tight junction biogenesis and neurite formation in neurons (27,31–33). Knockdown of TIAM1 in the PREX1 knockout PriGO9A cells reduced Lgl phosphorylation. Knockdown in PriGO8A cells, which express very low levels of TIAM1, had no effect on Lgl phosphorylation, consistent with the changes in PREX1 knockout PriGO9A cells being due to on-target effects of the siRNA. This second mechanism for Lgl1 phosphorylation is probably active in only a small subset of glioblastoma patients, based on comparative RNA-seq expression of PREX1 mRNA in 152 patients in the TCGA database.

As PREX1 requires PIP3 binding for its activation and the role of phosphorylation in regulating Lgl is well established, the experiments described here establish a new role for PREX1 in linking aberrant PI 3-kinase pathway signalling in glioblastoma to the disruption of normal polarity pathway signalling. A subset of glioblastoma cells appear to have a redundant pathway for Lgl phosphorylation involving TIAM1. Full-length TIAM1 has complex and not fully understood mechanisms of activation, some of which may involve the PI 3-kinase pathway (34). In PriGO9A glioblastoma cells, TIAM1 overexpression mainly involves shorter isoforms of TIAM1; these are probably amino-terminal truncated isoforms generated by alternate splicing. Amino terminal truncation of Tiam1 results in its stabilization and constitutive activation (35). TIAM1 may therefore bypass the need for aberrant PI 3-kinase pathway signalling to promote Lgl phosphorylation in these cells. While PREX1 knockout affected Lgl phosphorylation in one of the two patient-derived glioblastoma cells, motility was impaired in both, suggesting a non-redundant role for PREX1 in glioblastoma cell motility. Lgl phosphorylation, while necessary for motility (14), does not appear to be sufficient. Marei *et al*. (36) have shown that PREX1, but not TIAM1, binds the actin remodelling protein FLII and that this is necessary for Rac-dependant cell migration in NIH3T3 cells; this is a likely candidate for a second PREX1-dependent signalling event that is required for glioblastoma motility.

## METHODS

### Antibodies

The following antibodies were used: PREX1 (D808D) rabbit monoclonal, doublecortin rabbit polyclonal antibody, LLGL1 (D2B5A) rabbit monoclonal antibody; phospho-PKC Substrate Motif ([R/K)XpSX(R/K)] MultiMab™ rabbit monoclonal antibody mix; TIAM1 rabbit polyclonal antibody; doublecortin rabbit polyclonal antibody, all from Cell Signaling Technology (Danvers, MA, USA); GAPDH mouse monoclonal from Abcam (Cambridge, MA, USA); Flag M2 mouse monoclonal antibody from Sigma-Aldrich (Oakville ON, Canada).

### Cell culture

Glioblastoma cells were isolated from patients undergoing first surgical tumor resection at The Ottawa Hospital as described previously (13). Cells were grown as monolayers on tissue culture plates coated with laminin (Sigma-Aldrich, Oakville ON, Canada) using neurobasal A medium with B27 and N2 supplements and EGF and FGF2 (all from Thermo Fisher Scientific Inc., Waltham MA, USA). Cells were incubated in 5% O_2_ and 5% CO_2_ at 37°C.

### Generation of PREX1 knockout glioblastoma cells

Glioblastoma PREX1 knockout cells were generated by electroporation of preformed crRNA/tracrRNA/Cas9 complexes. For the crRNA:tracrRNA duplex, a crRNA sequence targeting exon 2 of *PREX1* (CGTTCTGCCGGATGCGATGC) was used (Dharmacon, Lafayette CO, USA). This was combined with tracrRNA-ATTO 550 (IDT DNA Technologies, Skokie, Ill, USA) to a final duplex concentration of 44μM to form the complete guide RNA complex. The complex was then heated at 95°C for 5 minutes. To form the Cas9 solution, for each well, 0.3μl of 61μM Cas9 nuclease stock solution (IDT DNA Technologies, Skokie, Ill, USA) was combined with 0.2μl of Resuspension Buffer R (IDT DNA Technologies, Skokie, Ill, USA). To form the crRNA:tracrRNA:Cas9 complex, for each well, 0.5μl of crRNA:tracrRNA complex was combined with 0.5μl of diluted Cas9 nuclease and incubated for 20 minutes at room temperature. PriGO8A cells, grown to 70-80% confluence, were then resuspended in Resuspension Buffer R to 500,000 cells per well. For each well, the following complex was prepared: 1μl crRNA:tracrRNA:Cas9 complex, 9μl cell suspension, and 2μl of 10 μM electroporation enhancer (Alt-R Cas9 Electroporation Enhancer, IDT DNA Technologies). To prepare the Neon Transfection System for electroporation, the Neon Tube was filled with 3ml of Electrolytic Buffer and inserted into the Neon Pipette Station. A Neon Tip was inserted into the Neon Pipette and 10μl of the 12μl solution available for each well was drawn into the tip. The Neon Pipette and Tip were inserted into the Pipette Station. The following electroporation parameters were used: 1050V, 30ms, and 2 pulses. After electroporation, cells were immediately plated on a laminin-coated 6-well plate. ATTO 550 fluorescence was verified 24 hours later by fluorescence microscopy. Two weeks after the first round of electroporation, a second round was performed on the same cells with the same experimental set-up and electroporation parameters.

### Cloning by limiting dilution

Cells were diluted to a concentration of approximately one cell per 100 uL and plated in 96 well plates. Growth in 96 well plates was done using a 50:50 mixture of regular media and conditioned media from 48 h cultures of untreated PriGO8A cells.

### TIDE assays

Genomic DNA was isolated using the Bio Basic All-4-One Genomic DNA MiniPrep Kit flowing the manufacturer’s protocol including the RNase treatment (Bio Basic Inc., Markham ON, Canada). The region around the CRISPR PREX1 target site was PCR-amplified using the primer pair DPREX3F (5’-GCACAGAGGGAAAGTCTCGG-3’) and DPREX3R (5’-GCTGCTCCAGTGTGTTTAAGG-3’). Sanger sequencing of PCR products was performed and sequence data was analyzed using TIDE software (https://tide.nki.nl/) (29).

### Western blotting

Western blotting was done as described previously (13). Blots were stained with amido black prior to probing with antibody, to confirm even transfer of proteins. Blots were probed with antibody to GAPDH as an additional loading control. Blots were imaged using a BioRad ChemiDoc Imaging System (BioRad, Mississauga ON, Canada).

### Immunofluorescence

Immunofluorescence was done as described previously (37).

### TERT promoter mutation analysis

The region of genomic DNA containing the TERT promoter region was PCR-amplified as described(38) and PCR products were Sanger sequenced as above.

### Live cell videomicroscopy

Cells were plated on laminin-coated Bioptechs delta-T dishes (Butler, PA, USA). For the duration of video acquisition, cells were maintained in a sealed chamber at 37°C and 5% CO_2_. Phase contrast images were taken at ten-minute intervals for 90 min total. Images were acquired with the 10X objective of a Zeiss Axiovert 200M microscope equipped with an AxioCam HRm CCD camera (Zeiss, Göttingen, Germany). Motility was quantitated using the MtrackJ plugin(39) in ImageJ software (National Institutes of Health, Bethesda, Maryland, USA) as described previously(14).

### Lgl1 phosphorylation assay

Lentiviral vectors expressing human flag-tagged Lgl1 and Lgl1(3SA) were described previously (13). Lentiviral particles were generated as described previously(13), concentrated using Lenti-X Concentrator (Takara, Mountain View CA, USA) and resuspended in neurobasal A medium supplemented as above. Phosphorylation of Lgl1 was detected by Western blotting with phospho-PKC Substrate Motif antibody mix; the blot was then stripped and re-probed with Lgl1 antibody. Non-transduced cells and cells transduced with LLGL1(3SA) were used as negative controls in all assays. *RNA-Seq:* Total RNA was isolated using the GE Illustra RNA Spin Kit (Thermo Fisher Scientific, Ottawa, ON, Canada) according to kit protocol including DNase treatment. RNA was eluted 2x with 40 uL of RNAse free water and stored at −80C. RNA concentration was assayed using a Nanodrop 1000 (Thermo Fisher Scientific, Ottawa, ON, Canada) and diluted to be within range of RNA seq requirements. (approx. 100 ng/uL). RNA-seq libraries were generated from 250 ng of total RNA. The NEBNext Poly(A) Magnetic Isolation Module and cDNA synthesis was achieved with the NEBNext RNA First Strand Synthesis and NEBNext Ultra directional RNA Second Strand Synthesis Modules (New England BioLabs, Ipswich, Ma, USA). Libraries were prepared using the NEBNext Ultra II DNA Library Prep Kit for Illumina (New England BioLabs, Ipswich, MA, USA). Paired-end 100bp reads were performed on an Illumina HiSeq4000. Pseudo alignment and transcript quantification were performed with Kallisto (40) and differential expression was determined using Sleuth (41). Gene ontology analysis was performed using Enrichr (42,43).

### Lentiviral vector constructs

A plasmid with cDNA for full-length human PREX1 was obtained from Dr. Heidi Welch (Babraham Institute, Cambridge, UK). To make a lentiviral vector expressing the DH-PH domain of PREX1, the primers 5’-GGATCCATGGAGGCGCCC AGCGGCAGC-3’ and 5’-CATCTTTGTAATCGCCCATGACGTAGGCATCACGCTC-3’ were used to amplify the region coding for PREX1 DHPH domains. A second round of PCR were then done with the same 5’ primer and the 3’ primer 5’-GAATTCTCATTTGTCGTCATCATCTTTGTAATCGCCCATG-3’ to add codons for a carboxy terminal Flag tag and a stop codon. The final PCR product was subcloned into pMiniT 2.0 vector (New England BioLabs, Whitby ON, Canada) and fully sequenced. cDNAs with the correct sequence were then subcloned into the doxycycline-inducible lentiviral vector pLVX-Tight-puro (Clontech, Mountain View CA, USA) using BamH1 and EcoR1 restriction sites. Lentiviral particles were generated as described above.

### siRNA knockdown

SMARTpool siGENOME TIAM1siRNA and siGENOME human TIAM1 (7074) siRNA 5’ CAUUCAAUCCUGCGUGAUA 3’ from Dharmacon (Lafayette, CO, USA) were used at final concentrations of 20 nM to knock down TIAM1 expression, as described previously(17).

### Statistics

Statistical analyses were done using SigmaPlot12 software. Details of specific tests used are given in the figure legends.

## DATA AVAILABILITY

All data are within the manuscript or available upon request. RNA-Seq data will be deposited to the Gene Expression Omnibus database.

## SUPPORTING INFORMATION

*Video S1.* PriGO8A cells. All videos were recorded as described in Methods.

*Video S2.* 8A/clone 4 cells.

*Video S3.* 8A/clone 6 cells.

*Video S4.* 8A/clone 18 cells.

*Video S5*. 8A/clone 4/iPREX1 no doxycycline

*Video S6.* 8A/clone 4/iPREX1 plus doxycycline

*Video S7.* PriGO9A

*Video S8.* 9A/clone 18 cells.

*Video S9*. 9A/clone 4/iPREX1 no doxycycline

*Video S10.* 9A/clone 4/iPREX1 plus doxycycline

